# BiHiTo: Biomolecular Hierarchy-inspired Tokenization

**DOI:** 10.64898/2026.01.23.701302

**Authors:** Ruochong Zheng, Yutian Liu, Yian Zhao, Zhiwei Nie, Xuehan Hou, Chang Liu, Siwei Ma, Youdong Mao, Jie Chen

## Abstract

Three-dimensional atomic arrangements of biomolecules are key to demystifying biological functions. The rapid expansion of accessible structural data, driven by advances in AI for science, highlights the critical challenge of efficiently modeling large-scale biomolecular structures, which are high-dimensional systems shaped by biological assembly principles. To address this, we introduce **BiHiTo**, a multi-level **Bi**omolecular **Hi**erarchy-inspired **To**kenizer that intrinsically mimics natural biological assembly hierarchies. Specifically, we design a multi-codebook quantizer that mirrors the natural hierarchy of biomolecular structure, enabling simultaneous capture of representations spanning atomic motifs to global conformational variations. This hierarchical alignment markedly improves the biological interpretability and reconstruction fidelity of biomolecular structure.Extensive experiments demonstrate that BiHiTo delivers state-of-the-art performance and robust generalization across molecular dynamics trajectories and macromolecular complexes, facilitating advances in structure generation and dynamic conformation exploration. In the reconstruction of the CASP14 and OOD test set FastFolding protein multi-conformation data, our method achieves a **17%** and **51%** reduction in RMSD compared to Bio2Token, respectively.

## Introduction

Biomolecular structures dictate the conformational landscapes and functional repertoires of macromolecules. High-resolution experimental techniques, most notably X-ray crystallography and cryo-electron microscopy, routinely resolve atomic coordinates with sub-angstrom precision. The resulting structural coordinates are archived in community repositories such as the Protein Data Bank (PDB) (wwp 2019). Complementarily, recent advances exemplified by AlphaFold2 (Jumper et al. 2021) have generated proteome-scale structural models approaching experimental accuracy, thereby expanding the accessible structural universe by several orders of magnitude. The concomitant expansion of high-quality structural data now enables systematic, datadriven investigations of macromolecular interaction networks (Abramson et al. 2024; Mirdita et al. 2022; Fang et al. 2025), catalyzing advances across fundamental biology and diverse biotechnology applications.

However, the structural complexity inherent in biomolecules presents a substantial challenge for computational modeling. Current structural generation methods, including diffusion models and autoregressive frameworks (Yim et al. 2023; Shin et al. 2021), often grapple with exponentially increasing computational complexity when simulating atomic-level interactions within large-scale biomolecular systems. This highlights the pressing necessity of developing all-atom encoding/decoding techniques.

Recent advances have made strides in the biomolecular structure tokenization. Several methods focus on small molecules (Zhou et al. 2023; Li et al. 2024; Gao et al. 2024). In the context of proteins, FoldSeek (Van Kempen et al. 2024) converts protein shapes into 1D sequences using structural alphabets, facilitating rapid comparisons. ESM-3 (Hayes et al. 2025) trains transformer models to encode protein backbones. Most of these approaches either focus on backbone atoms or are limited to training on short chains. More recently, unified biomolecular VAEs such as Bio2Token (Liu et al. 2025) treat biomolecules purely as point clouds to achieve uniformity. However, it ignores the inherent hierarchical priors of biological molecules, making training more difficult and hindering generalization.

Rather than treating biomolecules as generic 3D point cloud data, we recognize that they possess an inherent multi-level organizational structure. Taking proteins as an example, their three-dimensional conformations arise from the hierarchical assembly of thousands of atoms, which form secondary structural elements such as alpha-helices and beta-sheets that further organize into functional supramolecular complexes, *cf*. Fig. 1. This natural hierarchy can serve as structural priors for biomolecule structure encoding.

**Figure 1.**
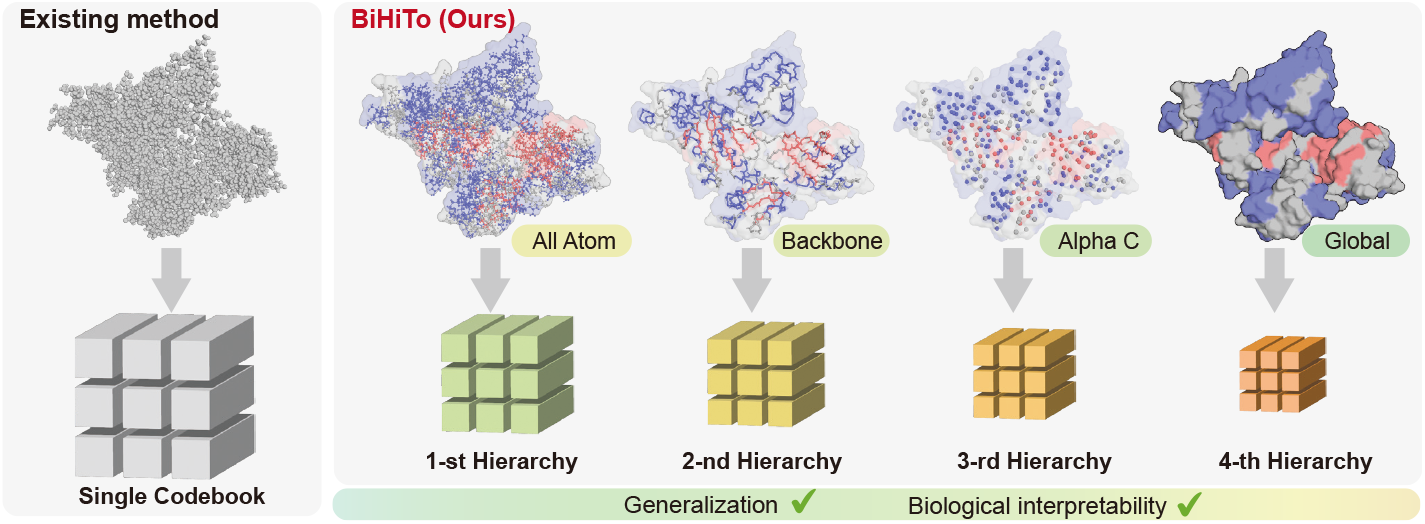
Biomolecular tokenization approaches. (Left) Conventional methods treat biomolecules as homogeneous point clouds with a single codebook. (Right) Our BiHiTo framework explicitly models biological hierarchy through multi-resolution representations, preserving structural priors for superior generalization and interpretability.

Building upon this, we introduce **BiHiTo**, a multi-level **Bi**omolecular **Hi**erarchy-inspired **To**kenizer that intrinsically mimics natural biological assembly hierarchies. Specifically, we design a multi-codebook quantizer that decomposes biomacromolecules into five biologically structured quantization levels: global topology, sparse sampling of *α*-carbon/C3^*′*^ intervals, full *α*-carbon/C3^*′*^ backbones, complete backbone atoms, and full-atom resolution.

Through this multi-level design, the quantizer explicitly encodes the intrinsic structural hierarchy of biomolecules, enabling simultaneous capture of representations spanning atomic motifs to global conformational variations. Extensive experiments demonstrate that the proposed BiHiTo significantly outperforms existing VQ-VAEs (Van Den Oord, Vinyals et al. 2017) for complex structures across all metrics, including reconstruction accuracy and stereochemical validity. BiHiTo also enhances the modeling efficacy and generalization capability, achieving a 25% lower RMSD on the RNA3DB test set and a 51% lower RMSD on the out-of-distribution FastFolding protein multi-conformation dataset than Bio2Token (Liu et al. 2025).

In summary, our main contributions are as follows:

- We design a multi-level biomolecular tokenizer based on the biomolecular structure hierarchical assembly prior, enabling simultaneous capture of representations spanning atomic motifs to global conformational variations.
- We employ a multi-codebook quantizer to model different natural hierarchies of biomolecules separately, enhancing model generalization while efficiently processing macromolecular complexes.
- Extensive experiments demonstrate that the proposed method achieves state-of-the-art performance in reconstruction accuracy and stereochemical validity.

## Related Work

### 3D Point Clouds Tokenization

Point clouds, as a 3D data format, present unique challenges for deep learning due to their inherent sparsity, disordered arrangement, and irregular structure. Researchers have developed various methods to convert unstructured point cloud data into ordered token representations. These tokenization approaches enable more efficient processing by neural networks, especially transformer-based architectures. For instance, PointContrast (Xie et al. 2020) and DepthContrast (Chhipa et al. 2022) both establish instance discrimination frameworks through contrastive learning, where the former aligns features of identical points across multi-view observations while the latter extends this paradigm to depth map augmentations for enhanced 3D representation learning. More recently, Point-BERT (Yu et al. 2022) adopts a BERT-like masked pre-training paradigm, and groups point clouds into local patches through farthest point sampling (FPS) (Eldar et al. 1997) for tokenization. Point-MAE (Pang et al. 2022) maintains the FPS-based patch partitioning scheme but transitions to an MAE-style framework. Point-M2AE (Zhang et al. 2022) further employs a hierarchical self-supervised learning to enhance global-to-local reconstruction. Current point cloud tokenization approaches show limited applicability to biomolecular structures, owing to the stronger intrinsic order inherent in biological systems. The geometric regularities of biomolecules, such as precise bond lengths, angular constraints, and torsional preferences, demand explicit structural modeling, a capability that point cloud methods inherently lack.

### Biomolecules Structure Tokenization

Current tokenization methods for biomolecular structures largely focus on single molecule types. For small molecules, UniMol (Zhou et al. 2023) tokenizes atoms individually with pairwise structural features; Geo2Seq (Li et al. 2024) encodes 3D geometries into SE(3)-invariant sequences; and MolStructTok (Gao et al. 2024) uses spherical line notation with VQ-VAE discretization.

Proteins, as large linear polymers with sequential order, offer natural advantages for structural discretization. Fold-Seek (Van Kempen et al. 2024) pioneered converting 3D backbone structures into 1D sequences via the ‘3Di’ alphabet. ProTokens (Lin et al. 2023) introduces an unsupervised framework integrating structure prediction and inverse folding to tokenize protein backbone structures into compact, amino-acid-like discrete representations. ESM-3 (Hayes et al. 2025) applies a transformer-based VQ-VAE with geometric attention, while InstaDeep (Gaujac et al. 2024) and FoldToken4 (Gao, Tan, and Li 2024) employ GNN-based vector quantization. Bio2Token (Liu et al. 2025) further introduces Mamba-based quantized auto-encoders for all-atom structures without SE(3) constraints.

In this work, we extend structure VQ-VAE to incorporate multiscale tokens reflecting the natural hierarchical organization of biomolecules, enabling high-fidelity reconstruction and biologically interpretable encoding.

## Method

In this section, we elaborate on the proposed method, which comprises three fundamental components: an encoder, a quantizer, and a decoder.

Next, we start by introducing the Mamba-based model architecture to ensure efficient and robust feature extraction. Then, we outline the proposed natural hierarchical quantization to achieve multilevel tokenization of biomolecular structure, which significantly enhances the model’s ability to capture and represent complex data patterns.

The details of our methodology are as follows.

### Model Architecture

In the encoder, a biomolecular structure with *N* heavy atoms is represented as a point cloud *X* ∈ ℝ^*N*×3^. This point cloud is atom-identity-agnostic and carries no residue or atom-type information. Our encoder and decoder employ Bidirectional Mamba (Gu and Dao 2023) Layers (BMLs), a neural module designed to capture bidirectional structural dependencies in biomolecular point clouds while preserving atomic locality.

Each BML employs a symmetric residual structure that leverages flip operations (Flip) and weight-shared Mamba blocks to establish bidirectional information flow. Given an input point cloud sequence *X* = [**x**_0_, **x**_1_, …, **x**_*N*−1_] ∈ ℝ^*N×d*^ of length *N*, processing occurs along two paths:

- Primary Path: *X* directly propagates to the final Add module (residual connection). This path preserves the original structural information and ensures stable gradient flow during backpropagation.
- Processing Path: *X* undergoes sequential transformations: Flip → Mamba Block → Flip. The Mamba Block processes the reversed sequence ℱ (*X*) with weight sharing (analogous to bidirectional RNNs (Zaremba, Sutskever, and Vinyals 2014)). This symmetric processing allows the model to capture long-range dependencies in both forward and reverse directions.

The output sequence **Y** is formulated as follows:

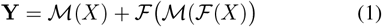

where *X* represents the input point cloud sequence and each component plays a specific role in bidirectional feature learning: ℱ denotes the flip operation that reverses the sequence order, enabling the model to process structural dependencies in both forward and reverse directions; ℳ corresponds to the Mamba Block’s transformation.

### Natural Hierarchical Quantization

As illustrated in Fig. 2, the NHQ component, as the core module, comprises three key aspects: multi-level downsampling leveraging biomolecular hierarchy priors, hierarchical quantization enabling expressive representations with computational efficiency, and feature fusion with reconstruction techniques incorporating upsampling and loss optimization. The details of these three parts are as follows.

**Figure 2.**
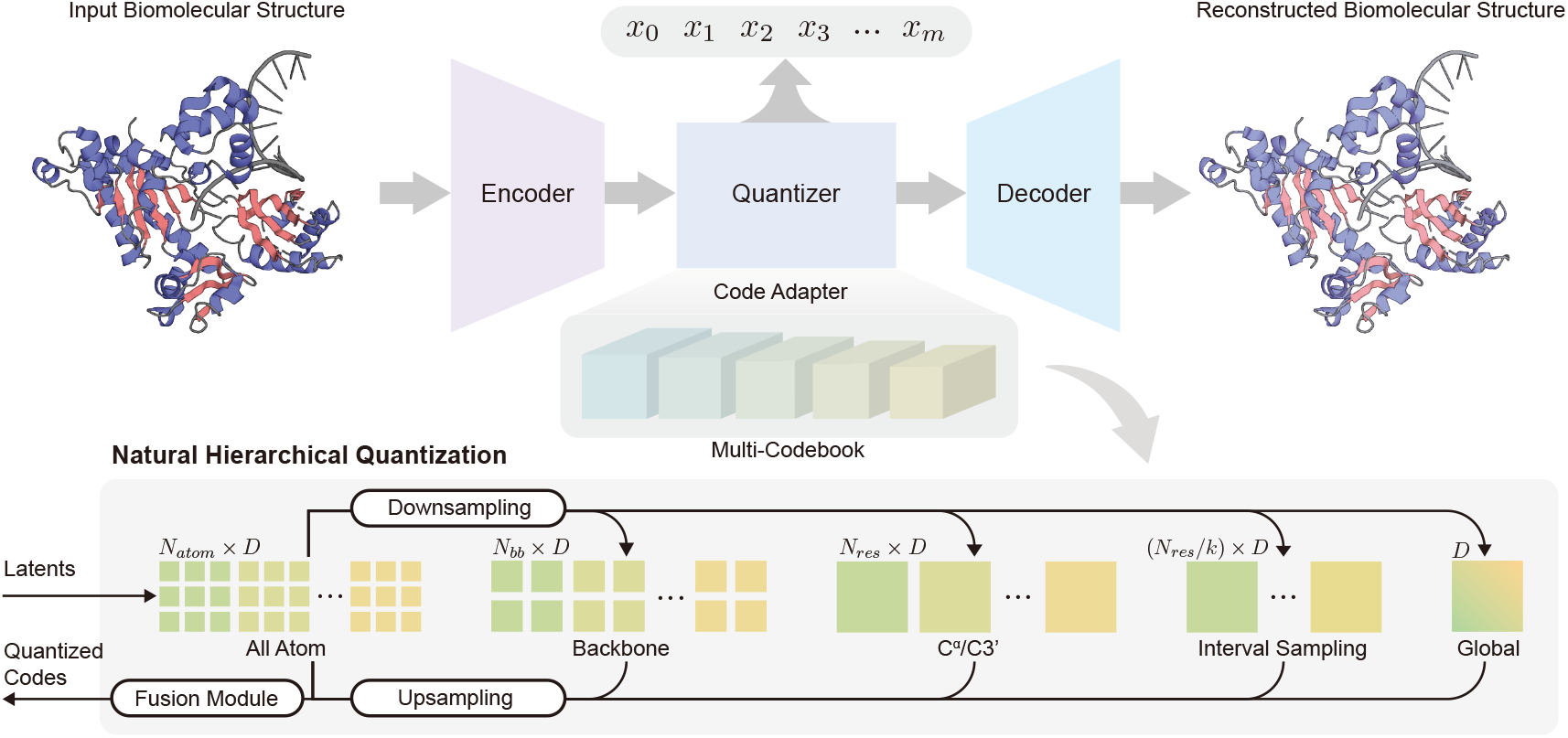
Natural Hierarchical Quantization Framework. Biomolecular embeddings from the encoder are processed through: (1) Hierarchical downsampling (Global topology → C_*α*_/C3^′^ Interval sampling → Full C_*α*_/C3^′^ → Backbone atoms → Full atoms); (2) Multi-level Finite Scalar Quantization (FSQ) with dedicated codebooks per hierarchy level; (3) Upsampling and feature fusion ; (4) High-fidelity reconstruction via Mamba-based decoder. This architecture preserves intrinsic biomolecular hierarchies while optimizing long-range dependency modeling.

#### Algorithm 1: Hierarchical Sampling Strategy

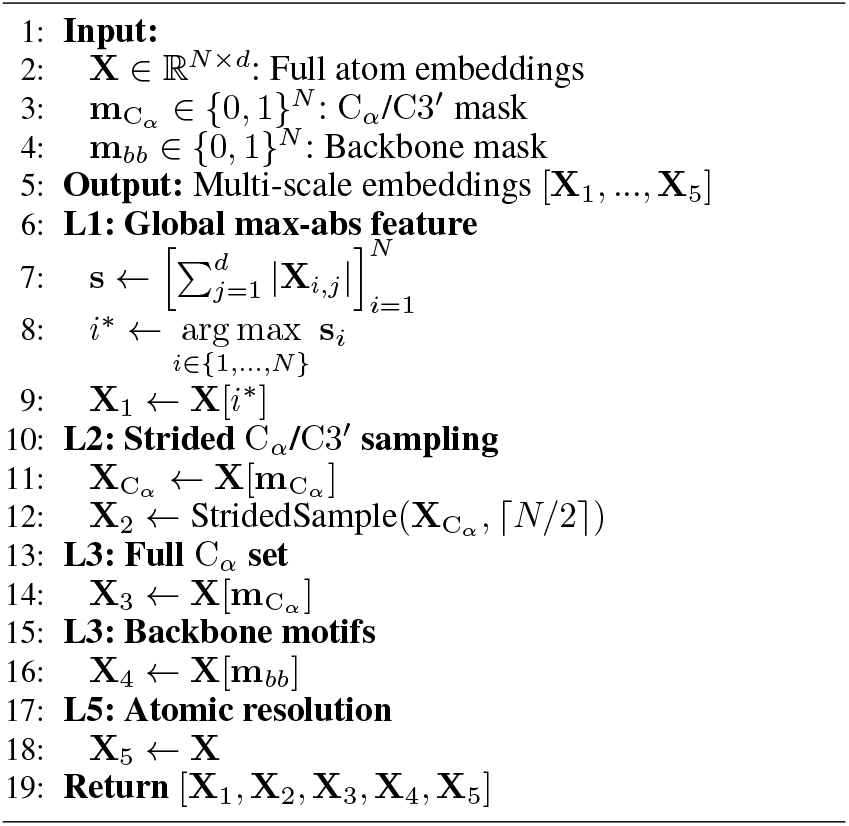

#### Natural Hierarchy

Conventional biomolecular reconstruction methods homogenize all atoms, disregarding the topological dominance of critical atoms (*e*.*g*., *α*-carbons, visualized in Fig. 1). To address this, we propose a physics-informed hierarchical embedding strategy. Through hierarchical sampling and residue-aware encoding, this strategy explicitly models atomic primacy relationships.

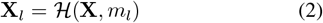

where **X**_*l*_ denotes the encoder-embedded molecular representation of level l, ℋ denotes the hierarchical sampling function and *m*_*l*_ specifies the sampling mask for each hierarchy.

The sampling rule dynamically adapts to both layer-specific physical properties and mask parameters *m*_*l*_. The hierarchical sampling strategy is formally defined as Alg. 1. The algorithm decomposes biomolecular representations into five distinct resolution levels:

- **Level 1 (L1: Global max-abs feature) s** represents the per-atom activation vector (sum of absolute feature values), *i*^∗^ denotes the index of the atom with maximum activation sum. This stage identifies the most biophysically significant atom as a topological anchor point, capturing the molecular “hotspot” with the highest combined feature activation. This provides a global reference for structural alignment and functional site identification.
- **Level 2 (L2: Strided** C_*α*_**/**C3^*′*^ **sampling)** 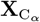 is the feature matrix of C_*α*_ (proteins) or C3^*′*^ (RNA) atoms, while **X**_2_ contains the uniformly subsampled C_*α*_/C3^*′*^ atoms. This level reduces the quantitative gap between global and C_*α*_/C3^*′*^ features, achieving a smooth transition between levels.
- **Level 3 (L3: Full** C_*α*_ **set) X**_3_ represents the complete set of C_*α*_/C3^*′*^ atoms. This stage maintains full backbone representation for precise modeling of residue-level conformation. Each C_*α*_/C3^*′*^ corresponds to one residue, directly encoding dihedral angles (*ϕ/ψ*) and secondary structure motifs, enabling accurate reconstruction of the polypeptide/nucleotide chain.
- **Level 4 (L4: Backbone motifs) X**_4_ contains the full backbone atom set (N, C_*α*_, C, O). This level enforces peptide bond rigidity constraints by including all backbone atoms, which is crucial for maintaining structural integrity and satisfying stereochemical constraints in protein folding.
- **Level 5 (L5: Atomic resolution) X**_5_ is the complete atomic embedding matrix. This final level preserves full atomic details for high-precision reconstruction of side-chain conformations, hydrogen-bond networks, and solvent interaction surfaces. It provides the finest granularity for atomic-level structural refinement, enabling accurate modeling of molecular interactions and binding sites.

#### Hierarchical Quantization

Within the Hierarchical Quantization (HQ) framework, our quantization system employs composite codebooks with exponential scaling. This structure enable compact high-fidelity representations while maintaining low search complexity. For each hierarchy level l, the quantization system is configured via a size vector 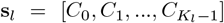 (where *K*_*l*_ denotes the sub-codebook count at hierarchy level *l, C*_*k*_ denotes dimension of sub-codebook 𝒞_*k*_ (number of learnable prototypes)). The system constructs an exponentially scaled quantization space via Cartesian product:

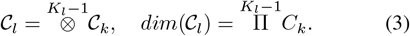

Each hierarchy level *l* maintains a dedicated codebook _*l*_, where the number of quantization steps |𝒞_*l*_| = *K*_*l*_ is pre-defined in the configuration file. Each quantization step *k* defines *C*_*k*_ fixed discrete states(learnable prototypes). Composite representations integrate multi-level features via:

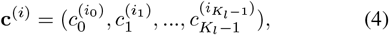

with index mapping:

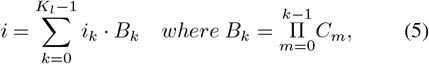

where base *B*_*k*_ denotes cumulative dimension scaling. Projection and Spatial Reorganization (Input Dimension Adaptation).

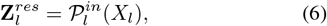

where 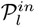 denotes the level-l input projection layer, transforming input **X**_**l**_ into residual features 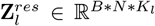 with batch size *B*, atomic count *N*, and quantization steps *K*_*l*_. This operation adapts the input dimensionality through linear projection and spatial reorganization to align with the hierarchical quantization structure.

Hierarchical Quantization Core can be represented as the following two steps:

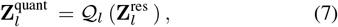

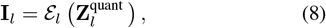

The quantization function 𝒬_*l*_ operates per quantization step with FSQ (Mentzer et al. 2023). The index encoding function ℰ_*l*_ calculates discrete indices through quantized projections.

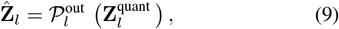

The output projection layer 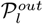 reconstructs features by mapping 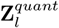 back to the original feature space, generating 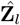 with dimensionality matching the input **X**_*l*_. This reconstruction optimizes feature fidelity through learnable linear transformations.

#### Up-Sampling

During feature reconstruction, we employ a direct block-wise repetition strategy for spatial resolution enhancement. Given the quantized downsampled feature matrix 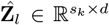 (*s*_*k*_: sampling points at level *k, d*: feature dimension), the upsampling ratio *r* is determined by atomic count *N* . The upsampling operation is defined as:

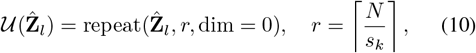

where 𝒰 denotes the upsampling operator and repeat( ) performs block-wise repetition along the spatial dimension (dimension 0).

#### Feature Fusion Mechanism

We integrate features from all five hierarchy levels through a concise fusion operation:

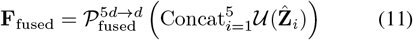

Here **F**_*fused*_ ∈ ℝ^*N×*5*d*^ represents the fused features, and the 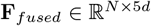 is a linear projection that restores the original feature dimension for decoder input. This design preserves hierarchical information while enabling adaptive weighting through learned projection parameters.

#### **Loss** Structural Alignment

Ground truth *X* and reconstructed 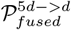 point clouds are aligned using the Umeyama-Kabsch algorithm. Loss functions are as follows:

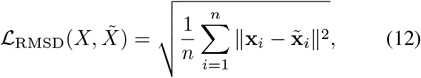

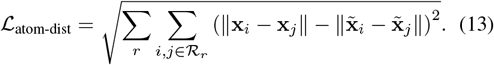

We use RMSD Loss and Inter-atomic Distance Loss that calculates the difference between the ground truth and reconstructed pairwise distances between each atom within a residue *r*. where *R*_*r*_ is the atomic index set within residue *r* (computed molecule-wide for small molecules). The total loss combines these components with equal weighting:

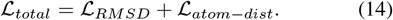

## Experiments

### Implementation Details

#### Datasets

We adopt the same training and testing data partitioning method as Bio2Token: (1) Protein training on CATH 4.2 (Ingraham et al. 2019) (18k structures; 40-500 residues) with topology-based splits, evaluated on CASP14/15 for reconstruction fidelity (up to 2,265 residues); (2) RNA training on RNA3DB (Szikszai et al. 2024) (≤10k atoms) with full-length testing; (3) Out-of-distribution generalization assessment using FastFolding’s 400k multi-conformation samples; (4)Small molecule training/testing on ∇^2^DFT (Khrabrov et al. 2024) dataset(1.9M molecules training, Test-Structure split for unseen 8-27 heavy atom evaluation).

#### Model and Training Configuration

For protein and RNA modeling, we implement a five-tier hierarchy quantization architecture: where level l (l=1,…,5) employs a codebook of size |𝒞_*l*_| = 4^*l*+1^, yielding configurations: L1:16, L2:64, L3:256, L4:1024, L5:4096 codewords. The encoder-decoder structure comprises 4 BML (encoder) and 6 BML (decoder) layers operating in ℝ^128^ For small molecule modeling, we exclusively utilize the L5 configuration (|𝒞_5_| = 4096) at the full-atom level.

The model is trained using the Adam optimizer with a polynomial learning rate schedule, initialized at 3 *×* 10^−4^. Training employs a batch size of 16 and a maximum sequence length of 10,000. All experiments are conducted on 8 NVIDIA RTX 3090 GPUs (24GB VRAM) for 216k steps (approximately 35 hours). L2 weight decay (*λ* = 0.01) is applied to prevent overfitting.

### Main Results

Protein structure reconstruction quality is evaluated via TM-score computed on C_*α*_ atoms (Zhang and Skolnick 2004) and Root Mean Square Deviation (RMSD). For RNA structures, TM-score is calculated based on C3^*′*^ atoms (Gong, Zhang, and Zhang 2019). TM-score measures local-to-global structural alignment (0: no similarity, 1: identical), while RMSD quantifies spatial deviation after aligning predicted and ground-truth point clouds using the Umeyama-Kabsch algorithm (Lawrence, Bernal, and Witzgall 2019). **Protein**. Based on the comparative analysis in Tab. 1, BiHiTo demonstrates state-of-the-art performance in protein structure prediction across diverse benchmark datasets. For the CATH4.2 test set, BiHiTo achieves an RMSD of 0.512Å, representing a 20.2% improvement over Bio2Token (0.642Å ) and significantly outperforming contrastive approaches such as ESM3. On the CASP14 benchmark, BiHiTo attains near-perfect structural accuracy with a TM-Score of 0.987 and RMSD of 0.515Å, comparable to the specialized protein model ProToken (TMScore 0.991, RMSD 1.229Å ).

**Table 1:**
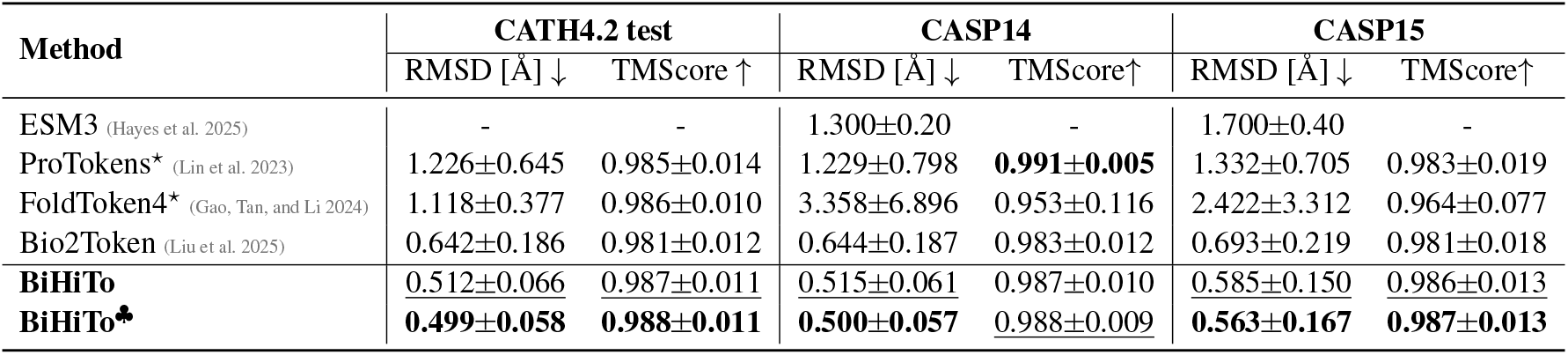
Summary of the best tokenizer models: RMSD and TM-Score between the ground truth structure and the reconstructed structure from the tokens. The results are presented in the format of “mean std”. ^*♣*^ denotes our method with the same amount of code (4096). ^⋆^ indicates that the model only reconstructs the protein backbone. **Bold** indicates the best performance and <u>underline</u> indicates the second best.

On the CASP15 benchmark featuring ultralong sequences (up to 2,265 residues), BiHiTo achieves an RMSD of 0.585Å . This is 75.8% lower than FoldToken4 (2.422Å ) and 15.6% better than Bio2Token (0.693Å ). This demonstrates BiHiTo’s strong generalization capability in the equilibrium protein structure.

#### RNA and Small Molecule

As shown in Tab. 2, the BiHiTo model significantly outperforms Bio2Token on the RNA3DB test set, achieving a 25.0% lower RMSD (0.578 vs. 0.771Å ) and an 8.7% higher TMScore (0.832 vs. 0.765). Notably, BiHiTo’s hierarchical quantization (L1–L5 sampling) effectively captures RNA structural features (*e*.*g*., backbone torsion angles). For long RNAs (*e*.*g*., Fig. 3 (b), PDB 8ZZJ, 4k atoms), BiHiTo reduces error accumulation (RMSD 0.53Å vs. Bio2Token’s 1.15Å )

**Table 2:**
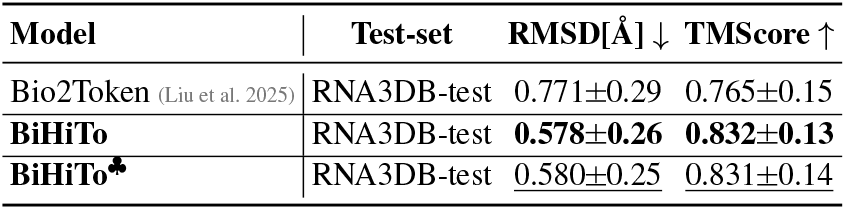
Comparison evaluation results of BiHiTo and Bio2Token on the RNA3DB test benchmark dataset and the Å small molecule benchmark dataset ∇^2^DFT/test-structure.

**Figure 3.**
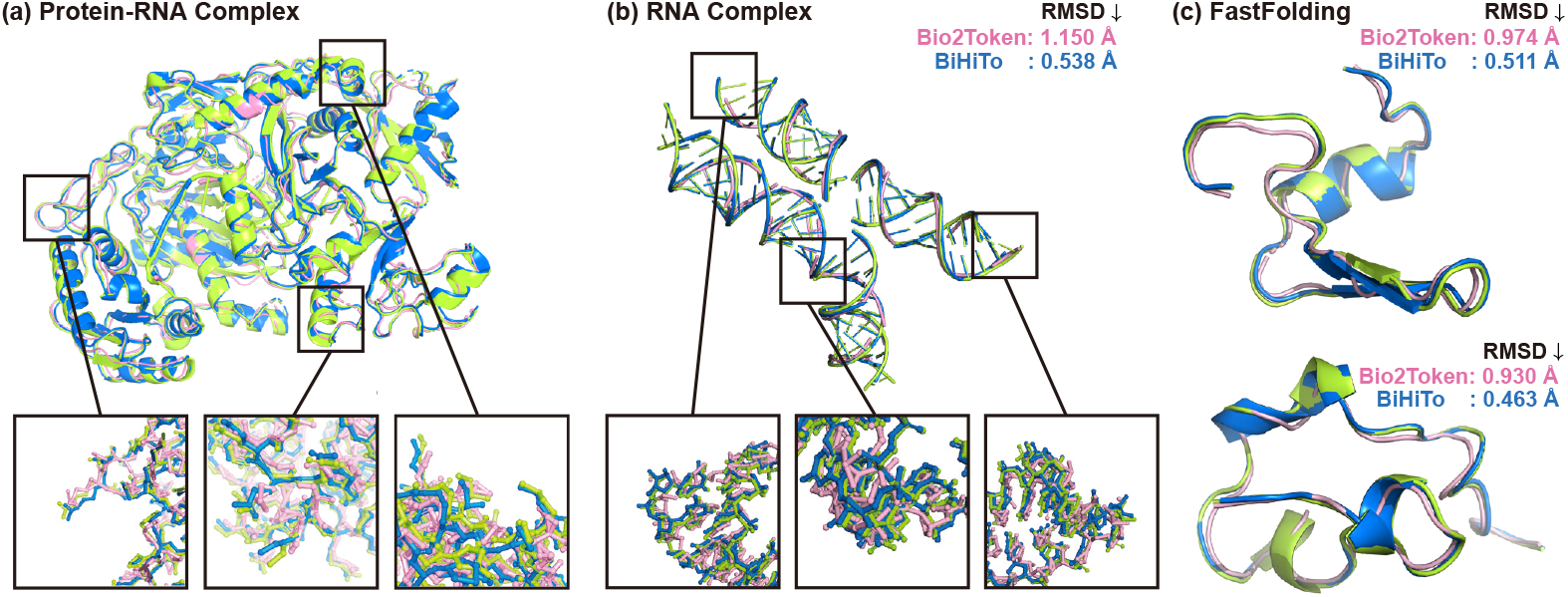
Reconstruction results of BiHiTo and Bio2Token in different scenarios, where green represents the ground truth, blue denotes BiHiTo results, and pink indicates Bio2Token results. (a) shows the reconstruction comparison of the two models on the complex structure 4W5O. (b) displays the structural reconstruction comparison of the models on the long-chain RNA 8JJZ. (c) presents the experimental results of the models reconstructing different conformations of the NTL9 structure, specifically frame0 and frame46500 from the FastFolding dataset.

In small molecule modeling, we exclusively employ L5 (full atomic level), identical to the Bio2Token architecture, bypassing unnecessary hierarchical complexity. Given the identical architecture and comparable codebook size to Bio2Token, the performance is nearly identical.

#### Complex

Notably, our model demonstrates outstanding performance on out-of-distribution data—specifically, molecular complexes. As shown in Fig. 3(a), BiHiTo achieves superior conformational reconstruction on the protein-RNA complex 4W5O, surpassing existing methods at atomic-level precision. Specifically, BiHiTo attains a RMSD of 0.666Å, representing a 46.5% reduction compared to Bio2Token’s 1.244Å . This result validates the unique advantage of our hierarchical quantization architecture in cross-molecular complex modeling.

### Generalization Experiments

#### Multi-Conformational Reconstruction

The experimental results in Tab. 3 demonstrate BiHiTo’s exceptional generalization capability on the FastFolding multi-conformational protein dataset. Crucially, this evaluation was conducted without any model training on this specific dataset. While Bio2Token achieved 0.965Å RMSD and 0.986 TM-Score, BiHiTo delivered breakthrough performance: RMSD decreased dramatically by 51.3% to 0.469Å while maintaining a high TM-Score of 0.996. This conclusively validates the cross-dataset generalization capacity of our architecture.

**Table 3:** Comparison of FastFolding protein multi-conformation dataset reconstruction results, tested on 400k conformations sampled every 100 frames from 40M protein conformations. **C***α***-dev** denotes the C_*α*_-C_*α*_ bond length deviation was calculated as the mean absolute deviation from the ideal value of 3.8Å .

| Methods | $C\alpha$-dev ↓ | RMSD ↓ | TM-Score ↑ |
| --- | --- | --- | --- |
| Bio2Token(Liu et al. 2025) | $0.455\pm0.05$ | $0.965\pm0.06$ | $0.986\pm0.006$ |
| **BiHiTo** | **$0.139\pm0.03$** | **$0.469\pm0.04$** | **$0.996\pm0.002$** |

To provide a more concrete illustration, we present in Fig. 3(c) a comparison between the predicted structures of the two models and the actual structures. This figure clearly demonstrates that our model exhibits superior predictive performance in both ordered and disordered regions of proteins. In the ordered regions, our predictions align almost perfectly with the actual structures, particularly excelling in the overlap accuracy of secondary structure elements such as *α*-helices and *β*-sheets, where as Bio2Token shows significant deviations in secondary structures. In disordered regions, our predictions better capture the conformational flexibility and dynamic properties of these segments. This robustly confirms the model’s strong generalization capability across diverse biomolecular systems.

### Ablation Study

#### Hierarchical Contribution Analysis

Through the formula sum(*W*_[*level*]_), we sum the absolute values of linear weights in the feature fusion layer to quantify the relative contribution of each level to the final representation. The experimental results in Fig. 4 clearly demonstrate the functional differentiation of different levels in structural modeling: L5 (full-atom level) exhibits the highest contribution, confirming the critical role of atomic-level details in precise position prediction. L3 (full *α*-carbon level) and L1 (global topology level) contribute comparably, revealing the synergistic importance of backbone conformation and global topology. L2 (sparse *α*-carbon sampling) and L4 (backbone atom level) show relatively lower contributions, reflecting their transitional nature.

**Figure 4.**
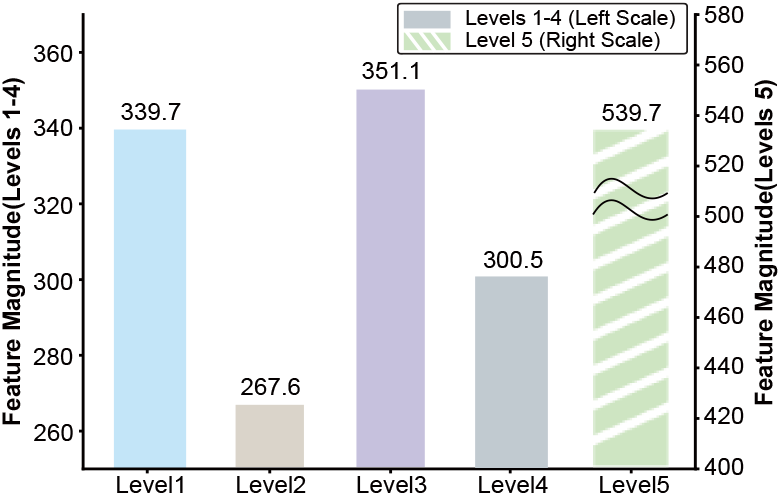
The contribution of each level is calculated by the formula sum(**W**_[level]_), where ∥**W**∥ *>* 0.01, which sums the absolute values of the effective weights at each level in the fusion layer.

To validate the structural necessity of each hierarchy, we conducted systematic ablation studies (*cf*. Tab. 4): Catastrophic impact of L3 removal: On out-of-distribution data like CASP15, removing L3 caused RMSD to increase by 16% with standard deviation expanding nearly 4-fold, indicating compromised generalization capability. Even on in-distribution RNA test sets, performance degraded, demonstrating that L3 ablation impairs both generalization and complex data performance, highlighting the irreplaceable role of *α*-carbon level in long-range folding accuracy.

Replaceability of L2: Removing L2 had a limited impact on conventional datasets (CATH, RNA3DB), but caused significant errors in CASP15 ultralong sequences (RMSD↑10.4%), confirming its auxiliary function in anchoring secondary structures within long sequences.

**Table 4:** Comparison of experimental results on the test set after removing different levels.

| BiHiTo/RMSD[$\text{\AA}$] | w all level | w/o level2 | w/o level3 |
| --- | --- | --- | --- |
| RNA3DB test | $0.578\pm0.26$ | $0.608\pm0.26$ | $0.622\pm0.29$ |
| CATH4.2 test | $0.512\pm0.06$ | $0.527\pm0.07$ | $0.528\pm0.07$ |
| CASP14 | $0.515\pm0.06$ | $0.537\pm0.10$ | $0.556\pm0.09$ |
| CASP15 | $0.585\pm0.15$ | $0.646\pm0.42$ | $0.682\pm0.59$ |

## Conclusion

By introducing hierarchical biological priors into biomolecular tokenization, BiHiTo achieves unified high-fidelity reconstruction across diverse biomolecules. Through multiscale quantization spanning global topology to atomic resolution, BiHiTo outperforms existing methods with 17% lower RMSD on CASP14 and 51% lower RMSD on out-of-distribution protein conformations. BiHiTo significantly enhances mainstream VQ-VAE architectures, demonstrating exceptional usability and robustness. Looking forward, BiHiTo enables efficient multi-scale biomolecular design to accelerate structural biology research.

## Acknowledgments

This work was supported in part by the Shenzhen Medical Research Funds in China (No. B2302037), National Natural Science Foundation of China (grant number 12125401), Natural Science Foundation of China (No. 61972217, 32071459, 62176249, 62006133, 62271465, 62406167), National Key Research and Development Program of China (grant number 2023YFF1204400 and 2023YFF1204401), and AI for Science (AI4S)-Preferred Program, Peking University Shenzhen Graduate School, China.

## Background

Three-dimensional atomic arrangements of biomolecules are key to demystifying biological functions. The rapid expansion of accessible structural data, driven by advances in AI for science, highlights the critical challenge of efficiently modeling large-scale biomolecular structures.

## Movation

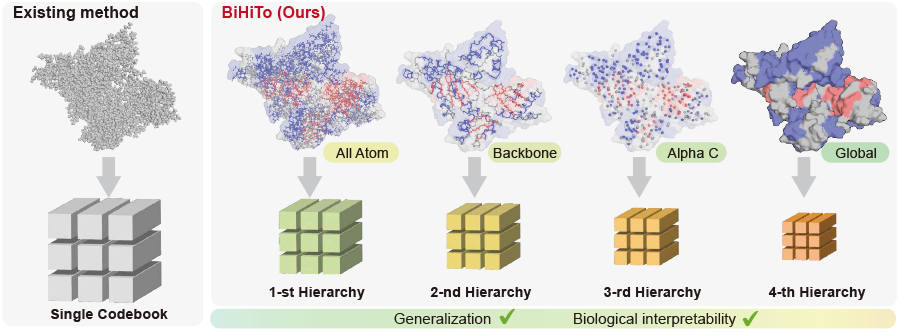

### Lack of structural priors

Previous SOTA it ignores the inherent hierarchical priors of biological molecules, making training more difficult and hindering generalization.

### Natural hierarchy

We recognize that they possess an inherent multi-level organizational structure. Taking proteins as an example, their three-dimensional conformations arise from the hierarchical assembly of thousands of atoms, which form **secondary structural** elements such as **alpha-helices** and **beta sheets** that further organize into functional supramolecular complexes.

## Contributions

- We design a **multi-level** biomolecular tokenizer **BiHiTo** based on the biomolecular structure hierarchical assembly prior, enabling simultaneous capture of representations spanning atomic motifs to global conformational variations.
- We employ a multi-codebook quantizer to model **different natural hierarchies** of biomolecules separately.
- In the reconstruction of the CASP14 and OOD test set FastFolding protein multi-conformation data, our method achieves a **17% and 51%** reduction in RMSD compared to the previous SOTA, respectively.

## Method

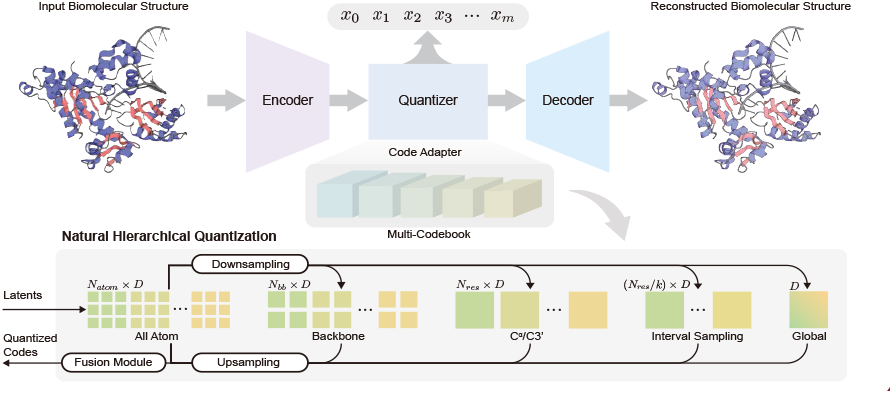

Natural Hierarchical Quantization Framework. Biomolecular embeddings from the encoder are processed through:

- Hierarchical downsampling (**Global topology** → **Cα/C3′** Interval sampling → **Full Cα/C3′** → **Backbone atoms** → **Full atoms**)
- **Multi-level** Finite Scalar Quantization (FSQ) with dedicated codebooks per hierarchy level
- Upsampling and feature fusion
- **High-fidelity** reconstruction via Mamba-based decoder

This architecture preserves intrinsic biomolecular hierarchies while optimizing long-range dependency modeling.

## Experiment

| Method | CATH4.2 test |  | CASP14 |  | CASP15 |  |
| --- | --- | --- | --- | --- | --- | --- |
|  | RMSD [Å] ↓ | TMScore ↑ | RMSD [Å] ↓ | TMScore ↑ | RMSD [Å] ↓ | TMScore ↑ |
| ESM3 (Hayes et al. 2025) | - | - | 1.300±0.20 | - | 1.700±0.40 | - |
| ProTokens* (Lin et al. 2023) | 1.226±0.645 | 0.985±0.014 | 1.229±0.798 | **0.991±0.005** | 1.332±0.705 | 0.983±0.019 |
| FoldToken4* (Gao, Tan, and Li 2024) | 1.118±0.377 | 0.986±0.010 | 3.358±6.896 | 0.953±0.116 | 2.422±3.312 | 0.964±0.077 |
| Bio2Token (Liu et al. 2025) | 0.642±0.186 | 0.981±0.012 | 0.644±0.187 | 0.983±0.012 | 0.693±0.219 | 0.981±0.018 |
| **BiHiTo** | <u>0.512±0.066</u> | <u>0.987±0.011</u> | <u>0.515±0.061</u> | 0.987±0.010 | <u>0.585±0.150</u> | <u>0.986±0.013</u> |
| **BiHiTo<sup>♣</sup>** | **0.499±0.058** | **0.988±0.011** | **0.500±0.057** | <u>0.988±0.009</u> | **0.563±0.167** | **0.987±0.013** |

On the CATH4.2 test set, the CASP14 benchmark, and the CASP15 benchmark, BiHiTo achieves reductions in RMSD of **20.2%, 22.3%**, and **15.6%** compared to the prior state-of-the-art, respectively, and demonstrates significant superiority over other competing methods.

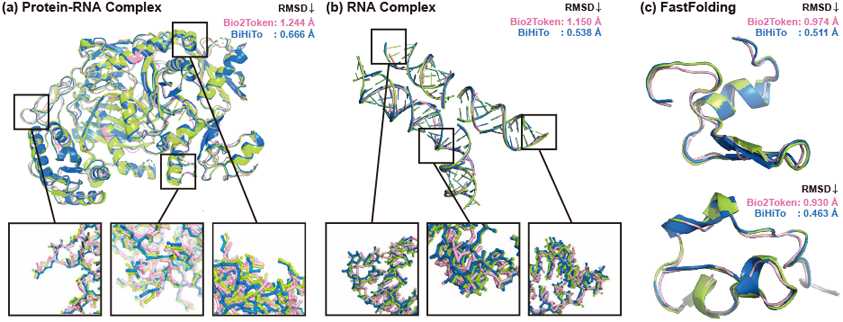

Our model also demonstrates outstanding performance on out-of-distribution data:

- **Complex Structure**: As shown in Figure (a) on the right, BiHiTo achieves a breakthrough in the conformational reconstruction of the protein–RNA complex 4W5O with atomic-level accuracy, significantly outperforming existing methods, which represents a **46.5%** reduction compared to Bio2Token.
- **Multi-conformation reconstruction**: Figure (c) on the right clearly demonstrates that our model achieves excellent predictive performance in both ordered and disordered regions of the protein, especially excelling in the overlap accuracy of secondary structure elements such as α-helices and β-sheets, with an RMSD of **0.511 Å**.

## Future Work

- Verify the effectiveness of BiHiTo on more downstream tasks.
- Apply BiHiTo to multi-scale biomolecular generation (similar to VAR structure) to accelerate the progress of structural biology research.

## Notes

### Competing Interest Statement

The authors have declared no competing interest.

