## Supplementary material for "BiHiTo: Biomolecular Hierarchy-inspired Tokenization"

### Supplementary material of "BiHiTo: Biomolecular Hierarchy-inspired Tokenization"

#### Open Source Statement

We will make our code public upon acceptance. The comprehensive visualization results and detailed implementation details in the paper and appendix fully demonstrate the reproducibility of our method. The training and testing code of the model, as well as the checkpoint, have all been submitted in the supplementary materials. The data can be downloaded from the link https://zenodo.org/records/15199953, and after placing it in the project directory, simply unzip it.

#### DataSet Detail

We used the data provided by Bio2Token for training and evaluation. Protein data utilizes the CATH 4.2 dataset for training/test, containing 18k protein structures from the PDB database with residue lengths ranging from 40 to 500 residues and heavy atom counts between 282 and 4,173. The dataset is partitioned according to the CATH topology classification defined by Ingraham et al., ensuring structural diversity between training and test sets. To evaluate generalization capability, we employ independent test sets CASP14 and CASP15 published post-CATH 4.2, which contain complex structures up to 2,265 residues (18,042 heavy atoms). For RNA data, the RNA3DB dataset is adopted, covering RNA structures in the PDB with nucleotide lengths spanning 2 to 4,450 nucleotides and heavy atom counts ranging from 42 to 95,518. To enhance training efficiency, sequence lengths were capped at $\leq$ 10,000 during training while full-length structures were included during testing. In addition, we utilized the FastFolding multi-conformation protein dataset as a test set to validate the model's generalization capability.testing 26 trajectories and 400K conformations across 12 proteins.We sampled every 100 frames across 40 million conformations from 26 trajectories of 12 proteins, ultimately obtaining 400,000 conformations. Generalization tests for BiHiTo and Bio2Token were then conducted on this dataset.

For small molecule data, we utilize the $\nabla^2 DFT$ dataset comprising 1.9 million molecules with 16 million conformations. Molecules contain 8–27 heavy atoms, and we employed the Test-Structure portion of this dataset for small molecule testing to evaluate unseen molecules and their conformations.

Copyright © 2026, Association for the Advancement of Artificial Intelligence (www.aaai.org). All rights reserved.

#### Case

As shown in Figure 1, BiHiTo demonstrates significant generalization performance improvements in multi-type biomolecule reconstruction tasks. In the multi-protein complex scenario (Figure a), BiHiTo reduces the average RMSD to 0.855Å, a 45.9% decrease compared to Bio2Token (Bio2Token: 1.583Å). The reconstruction accuracy improvement is even more pronounced in protein-RNA complexes (Figure b) (BiHiTo: 1.244Å vs Bio2Token: 0.675Å, RMSD=-45.7%). Most notably, for protein multi-conformation data with conformational diversity (Figure c), BiHiTo successfully reconstructs key structural domains such as -helices and -sheets (RMSD=0.390Å), achieving 60.6% higher accuracy than Bio2Token (Bio2Token: 0.990Å). 3D structure visualization further confirms that BiHiTo not only more accurately captures protein-nucleic acid interfaces but also maintains continuous conformations at complex topological junctions, enabling high-precision reconstruction in both ordered and disordered regions. This breakthrough overcomes the generalization bottleneck of traditional methods in cross-molecule-type reconstruction.

#### Quantization Detail

Hierarchical Quantization Core can be represented as the following two steps:

$$\mathbf{Z}_l^{quant} = \mathcal{Q}_l\left(\mathbf{Z}_l^{res}\right), \tag{1}$$

$$\mathbf{I}_l = \mathcal{E}_l\left(\mathbf{Z}_l^{quant}\right), \tag{2}$$

The quantization function $\mathcal{Q}_l$ operates per quantization step with FSQ. The index encoding function $\mathcal{E}_l$ calculates discrete indices through quantized projections.

$$\mathbf{Q}_l(\mathbf{Z}) = \frac{\tanh(z^{(k)} + \delta_k)\cdot \alpha_k - \beta_k}{\gamma_k} \quad k \in [0, K_l - 1] \tag{3}$$

where $\alpha_k = \frac{(C_k-1)(1-\epsilon)}{2}$, scales the dynamic range of feature values; $\epsilon$ is a small constant introduced for numerical stability. $\beta_k = \left\{ \begin{array}{ll} 0.5 & C_k \text{ is even} \\ 0 & C_k \text{ is odd} \end{array} \right.$, provides symmetry

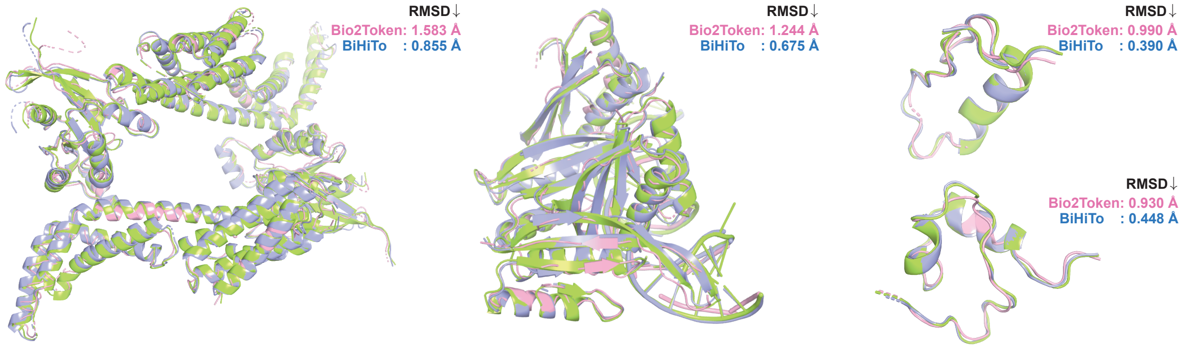


Figure 1: Reconstruction results of BiHiTo and Bio2Token in different scenarios, where green represents the ground truth, blue denotes BiHiTo results, and pink indicates Bio2Token results. (a) Comparison of the reconstruction of two models on the multi-protein complex 2H7V. (b) Comparison of structural reconstruction of protein-RNA complex 3WBM (c) The experimental results demonstrate the model's reconstruction of different conformations of the NTL9 and PRB structures, specifically frame3300 and frame2600 from the FastFolding dataset.

| **Test Set** | **FPS sampling (RMSD(time))** | **Interval sampling (RMSD(time))** | **Time increase percentage** |
|---|---|---|---|
| CATH4.2 test | 0.515 (13.29) | 0.512 (9.75) | 36.30% |
| CASP14 | 0.522(1.41) | 0.515(1.18) | 19.57% |
| CASP15 | 0.597740(2.49) | 0.585(1.98) | 25.96% |
| RNA3DB-test | 0.578(12.59) | 0.578(11.09) | 13.53% |

Table 1: The RMSD and time comparison of the model inferring 100 times on different test sets, when the sampling strategy of level2 is set to FPS sampling and interval sampling respectively.

compensation for even/odd centroids. $\delta_k = tan^{-1}(\beta_k/\alpha_k)$, shifts the hyperbolic tangent activation. $\gamma_k = \lfloor C_k/2 \rfloor$, normalizes quantization bins.

Index encoding $E_l$ computes discrete indices via:

$$\mathbf{E}_l(\hat{\mathbf{Z}}) = \sum_{k=0}^{K_l-1} \left[ \frac{\hat{\mathbf{Z}} \cdot \gamma_k - \beta_k}{\alpha_k} + 0.5 \right] \cdot \mathbf{B}_l^{(k)} \tag{4}$$

with basis vector $B_l = [1, \prod_{k=0}^{0} C_k, \cdots, \prod_{k=0}^{K_l-2} C_k]$. This encoding maps continuous quantized values to discrete indices, transforming feature space into hierarchical codebook

$$\hat{\mathbf{Z}}_l = \mathcal{P}_l^{out}\left(\mathbf{Z}_l^{quant}\right), \tag{5}$$

The output projection layer $\mathcal{P}_l^{out}$ reconstructs features by mapping $\mathbf{Z}_l^{quant}$ back to the original feature space, generating $\hat{\mathbf{Z}}_l$ with dimensionality matching the input $\mathbf{X}_l$. This reconstruction optimizes feature fidelity through learnable linear transformations.

#### FPS vs interval sampling

Although point clouds lack inherent hierarchical divisions, methods like FPS (Farthest Point Sampling) can still be used to establish levels. We therefore treat protein structures as point clouds and exclusively apply FPS sampling at Level 2, selecting $\sqrt{N_{C_\alpha}}$ alpha-carbons as the sampled representation for this hierarchy.

As shown in Table 1, replacing FPS sampling with our Alpha-Carbon Interval Sampling strategy in Level 2 significantly enhances computational efficiency while maintaining reconstruction accuracy. On the CATH test set, interval sampling improved RMSD from 0.515Å (FPS) to 0.512Å while reducing computation time by 36.30% (13.2967→9.7554 seconds). This advantage is more pronounced in CASP benchmarks: CASP14: RMSD decreased from 0.522Å to 0.515Å with 19.57% time reduction (1.4193→1.1870 sec); CASP15: RMSD improved from 0.598Å to 0.585Å with 25.96% time savings (2.4958→1.9813 sec). Notably, on the RNA3DB dataset, interval sampling achieved identical reconstruction accuracy (0.578Å) while still reducing time by 13.53% (12.5949→11.0937 sec).

The substantial time increase caused by FPS sampling – even when sampling only sparse points at Level 2 – highlights a critical limitation. Universal application of FPS across all hierarchies would further exacerbate computational costs. These results conclusively demonstrate that our biologically-informed hierarchical partitioning outperforms non-native point cloud partitioning methods in both effectiveness and computational efficiency.
